# CNOT1 mediates phosphorylation via Protein kinase A on the circadian clock

**DOI:** 10.1101/630897

**Authors:** Yunfeng Zhang, Haitang Qin, Yongjie Feng, Peng Gao, Yingbin Zhong, Yicheng Tan, Han Wang, Ying Xu, Joseph S. Takahashi, Guocun Huang

## Abstract

At the core of the mammalian circadian feedback loop, CLOCK (NPAS2)-BMAL1 is the positive element to activate transcription of downstream genes encoding the negative elements PERs and CRYs. Here we show that CNOT1 associates with both CLOCK and BMAL1, promotes their phosphorylation and increases their protein stability, and in turn inhibits the transcriptional activity of CLOCK-BMAL1. Expression of either CLOCK, BMAL1 or CNOT1 could interact with endogenous Protein Kinase A (PKA) as assessed by co-immunoprecipitation (Co-IP) and kinase assays. PKA could phosphorylate CLOCK and BMAL1 and this was promoted by CNOT1. Genetic deletion of *PKA-Cα* by CRISPR-Cas9 results in a longer period of the circadian rhythm; while overexpression of PKA-Cα induces a shorter period. Furthermore, we demonstrate that CNOT1 associates with CLOCK and BMAL1 in the mouse liver and promotes their phosphorylation. PER2, but not CRY2, is also a PKA target. Our results suggest that CNOT1 and PKA play a critical role in the mammalian circadian clock, revealing a conserved function in eukaryotic circadian regulations.

## Introduction

The circadian clock is an endogenous oscillator used to predict and adapt to the daily changes imposed by the 24 h light/dark cycle. At the molecular level, the core clock mechanism is composed of positive elements and negative elements to form a negative feedback loop conserved from bacteria to humans. In mammals, CLOCK-BMAL1 (or NPAS2-BMAL1) is the positive element and activates expression of its transcriptional repressors or negative elements, *Period* (*Per1* and *Per2*) and *Cryptochrome* (*Cry1* and *Cry2*) (1). A secondary loop is involved in RORα and REV-ERBα (NR1D1), which respectively activates or represses *BMAL1* transcription to stabilize and fine-tune the clockwork (2,3). There is also a third D-box loop involving the PAR-bZip transcriptional factors DBP, TEF and HLF, which are repressed by NFIL3 (E4BP4) (4). In addition, other feedback loops also include Dec1, Dec2 (5) and TRAP150 (6) as negative or positive regulators of CLOCK-BMAL1 respectively.

Post-translational modifications are important for stability, interaction and activity of clock proteins. For example, phosphorylation of CLOCK weakens its transactivation activity (7,8). SUMOylation is required for BMAL1 protein rhythmicity and normal clock function (9). O-GlcNAcylation of CLOCK, BMAL1, PER2 and dPER increases their stability and acts a metabolic signal (10–13). Ubiquitination of BMAL1 by the E3 ubiquitin ligase UBE3A promotes in its proteasomal degradation (14,15).

Interestingly, ubiquitination of CRYs by the F-box-type E3 ligase FBXL3 and FBXL21 has antagonizing actions on CRYs in different subcellular locations (16,17). Among those posttranslation modifications mentioned above, phosphorylation regulation has been widely identified and plays a key role in the circadian feedback loop (18).

Rhythmic phosphorylation of clock proteins by kinases and dephosphorylation by phosphatases are important for the generation and fine-tuning of circadian oscillations (7,19). Casein kinase I (CKI) is the most conserved kinase to regulate clock components and has been extensively studied in different organisms (18). Similar to CKI function, CK2 targets both PER2 and BMAL1 in mammals (20,21). Phosphorylation by GSK3 results in both CLOCK and CRY2 for proteasomal degradation (22,23); but PER2 phosphorylation by GSK3 promotes its nuclear entry (24). The protein kinase C-α (PKCα) and receptor for activated C kinase-1 (RACK1) rhythmically bind to the BMAL1 complex in the mouse liver, and RACK1 promotes BMAL1 phosphorylation by PKCα *in vitro* (25). PKCγ interacts and stabilizes with BMAL1 by reducing its ubiquitination (26). In contrast, both PKCα and PKCγ phosphorylate and activate CLOCK (27).

The cAMP-dependent PKA is implicated only in the specific output behavior of downstream of the clock and is not involved in the core clock in *Drosophila melanogaster* (28). However, recent data suggest that PKA stabilizes both PER and TIM in fly clock neurons (29,30), and PKA could phosphorylate TIM *in vitro* (31). In *Neurospora*, PKA is a kinase for stabilizing FRQ and WC proteins (32) and is also responsible for WC-independent transcription of the *frq* gene (33). PKA participates in the photic signaling pathway in mammals (34), however, whether PKA functions in the core clock proteins still remains unknown.

The Not1 protein is a subunit of the CCR4-NOT complex, which is a multisubunit protein complex that is conserved in eukaryotic organisms (35,36). It was first reported in yeast to act as an essential nuclear protein for negatively regulating transcription more than two decades ago (37). This protein was also identified in a large complex and was renamed from CDC39 to Not1 (<u>N</u>egative <u>o</u>n <u>T</u>ATA) (38). The CNOT (<u>C</u>CR4-<u>NOT</u>) complex proteins are also found in human cells (39,40). CNOT1 is the largest subunit and it is thought to be a scaffold for the complex, which plays many different cellular functions (35,36). We recently observed that Not1 stabilizes WC proteins and inhibits transcription of the clock gene *frq* in fungi (41), nevertheless little is known regarding the underlying mechanism.

Here we found that CNOT1 inhibited the transcriptional activity of CLOCK-BMAL1. The functional domain of CNOT1 in this process was located on the C-terminus, which possibly forms the “NOT module” including NOT2, NOT3, NOT4 and NOT5 proteins in yeast (42). Separate coexpression of CNOT1-C promotes phosphorylation and increases stability of both CLOCK and BMAL1. Overexpression of CNOT1-C, CLOCK and BMAL1 could, respectively, pull down endogenous PKA, and CNOT1-C promotes CLOCK binding to PKA. However, the *cnot1* knockdown decreases phosphorylation levels and protein amounts for both CLOCK and BMAL1. Phosphorylation mediated by CNOT1 was compromised if some potential PKA phosphorylation sites were mutated in the CLOCK and BMAL1, while at the same time these mutations increased transactivation of CLOCK-BMAL1. Transient transfection experiments showed that PKA could phosphorylate both CLOCK and BMAL1, but the PKA dead kinase could not. The deletion of *PKA-Cα* by CRISPR-Cas9 resulted in a phase advance and a long period; whilst overexpression of PKA-Cα displayed a phase delay and a short period. In CLOCK-HA transgenic mice, CNOT1 was co-immunoprecipitated with CLOCK-HA, promoting phosphorylation of both CLOCK and BMAL1 by PKA. PER2, but not CRY2, was also a PKA substrate, and CNOT1 and PKA promote PER2 phosphorylation and stability. Taken together, these data suggest that PKA phosphorylation mediated by CNOT1 plays an essential role in the mammalian circadian clock.

## Results

### CNOT1 suppresses transcriptional activity of CLOCK-BMAL1 by promoting its phosphorylation and increasing its stability

Given that Not1 is a conserved protein among different eukaryotic organisms and it associates and stabilizes WC proteins in fungi (41), we wondered whether it could play a similar role in the mammalian clock. Transient transfection assays using HEK293T cells showed that CNOT1 strongly inhibited the transcriptional activity of CLOCK-BMAL1 driving the *Per1* promoter (Fig. 1A), consistent with its function regulating the core clock *frq* gene in *Neurospora* (41). Not1 is a large protein containing more than two thousands amino acids and is composed of at least three protein-binding domains (42) in yeast. To test which domain of CNOT1 was essential for its suppression in mammals, three fragments were cloned, including CNOT1-N (amino acids 1-942), CNOT1-M (amino acids 882-1467) and CNOT1-C (amino acids 1407-2376). The reporter assay results showed that only the C-terminus was required for the CNOT1 function in this process (.1B). Overexpression of HA-tagged CNOT1-C with Flag-tagged CLOCK or Flag-tagged BMAL1 separately or together demonstrated interactions between CNOT1-C and CLOCK or BMAL1 (Fig. 1C). CNOT1-C also promoted accumulation of CLOCK or BMAL1 proteins respectively (Fig. 1D). Cycloheximide (CHX) treatment assays confirmed that CNOT1 stabilized both CLOCK and BMAL1 (Fig. 1E and F).

**FIGURE 1.**
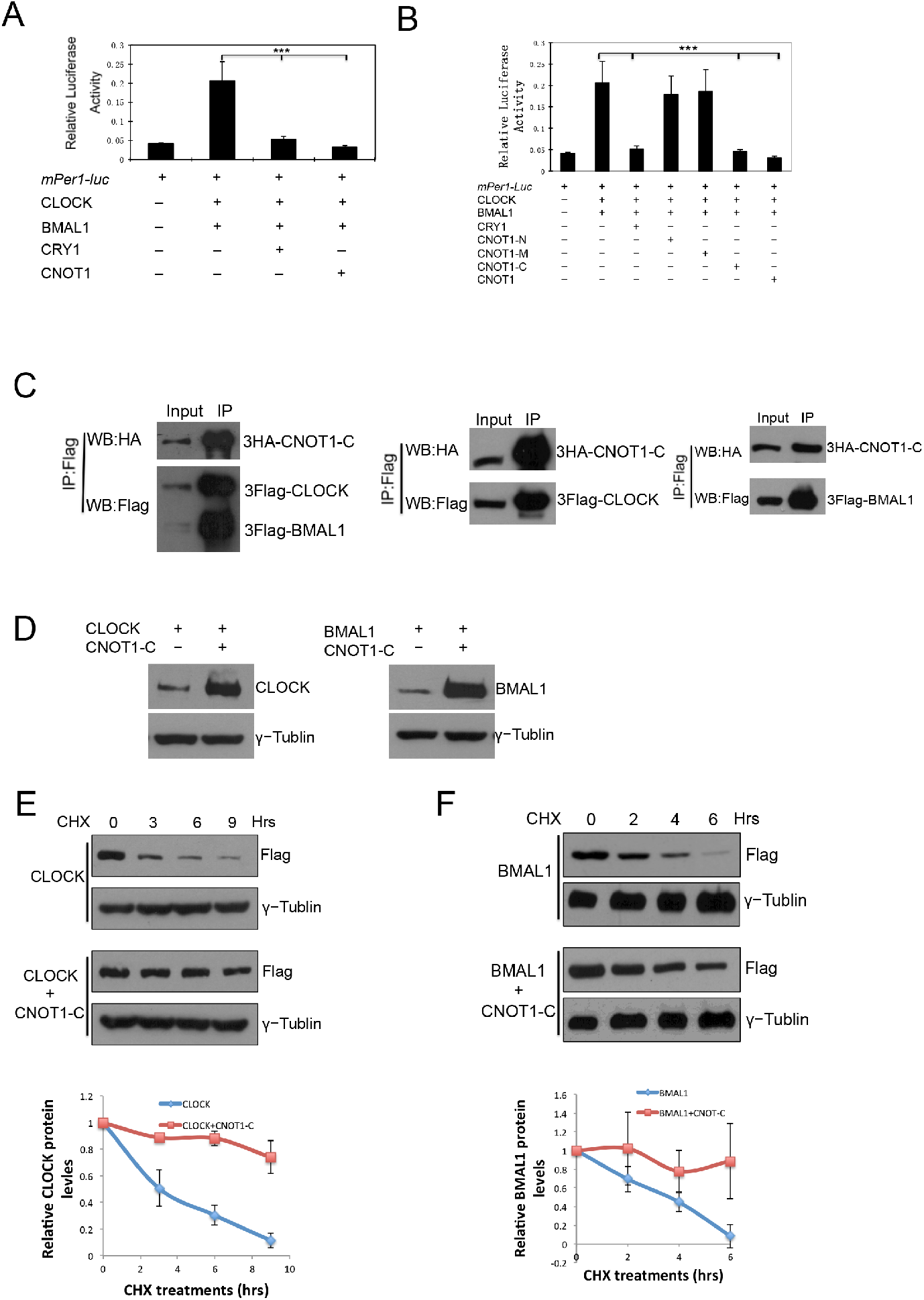
CNOT1 inhibits transcriptional activity of CLOCK-BMAL1 by interaction and stabilization of both CLOCK and BMAL1. (A) Effect of CNOT1 on transactivation by CLOCK-BMAL1 of a luciferase reporter gene driven by the *mPer1* promoter. Data are means (±SD). N=3. A two tailed Student’s *t*-test was performed, and significant differences are represented by black asterisks. (***) *P*<0.001. (B) The carboxyl-terminus of CNOT1 is important for its inhibition on CLOCK-BMAL1. (C) Interactions were shown between CNOT1 and CLOCK or BMAL1 in HEK293T cells. (D) CNOT1 stabilized both CLOCK and BMAL1 in HEK293T cells. (E and F) Western blot analysis showed that CNOT1 increased stability of CLOCK (E) and BMAL1 (F) after addition of CHX (20 μg/ml). Densitometric analyses of the results from three independent experiments were shown below.

### Phosphorylation of CLOCK and BMAL1 promoted by CNOT1 is regulated by PKA

In *Neurospora*, hyper-phosphorylated WC proteins are stable and have a weak DNA binding activity (43), and both WC-1 and WC-2 phosphorylation levels are compromised by the down-regulation of *not1* expression (41). To examine whether CNOT1 promotes phosphorylation of both CLOCK and BMAL1, a special gel (bisacrylamide:acrylamide=1:149) was employed to visualize high resolution of phosphorylation patterns (44). Hyper-phosphorylated bands of both CLOCK and BMAL1 were displayed when coexpressing CNOT1-C with CLOCK and BMAL1 respectively (Fig 2A). On the contrary, in the *cnot1* knockdown U2OS cells, both CLOCK and BMAL1 phosphorylation and amount decreased (Fig. 2B and 2C), suggesting CNOT1 has a same function on CLOCK and BMAL1 *in vitro* and *in vivo*. Then we asked which kinase is responsible for the specific phosphorylation promoted by CNOT1. Given that the Ccr4-Not complex components could regulate PKA activity in *S. cerevisia* (45) and PKA is a kinase for both WC-1 and WC-2 in *Neurospora* (32), we predicted that both CLOCK and BMAL1 could be targets of PKA. There are 16 and 11 potential PKA Phosphorylation sites among 167 and 100 Serine/Threonine of the CLOCK and BMAL1, respectively, based on phosphorylation prediction analysis. Four sites of CLOCK (S440, 441, 478, and 479) and two sites of BMAL1 (S42 and 246), which exhibit the highest probability of being phosphorylated by PKA, were mutated to alanines. The phosphorylation levels of both CLOCK and BMAL1 mutants were decreased compared to WT proteins when coexpressing with CNOT1 (Fig. 2D), suggesting that phosphorylation of CLOCK and BMAL1 promoted by CNOT1 could be regulated by PKA. As expected, the CLOCK/BMAL1 mutants showed higher transcriptional activity and less stability than their WT counterparts (Fig. 2E), similar to the data in fungi (43). PKA-Cα (PKA catalytic subunit alpha) dramatically inhibited transcriptional activity of CLOCK-BMAL1 (Fig 2F). Taken together, our results indicated that PKA phosphorylation inhibited CLOCK-BMAL1-mediated transcriptional activity and increased their protein stability.

**FIGURE 2.**
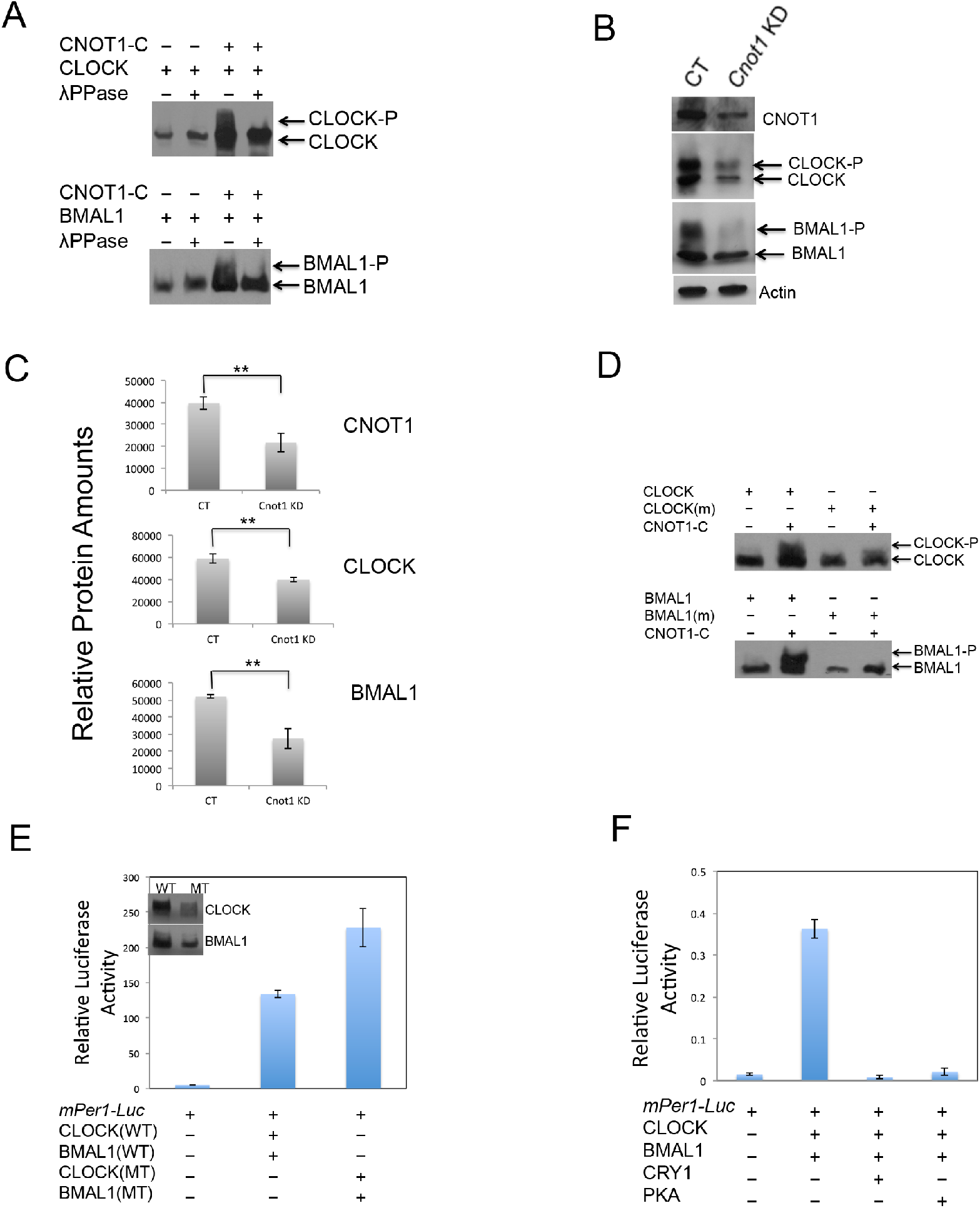
CNOT1 promotes phosphorylation and inhibits transcription of CLOCK-BMAL1. (A) Coexpressing CNOT1 induced phosphorylation of CLOCK and BMAL1 respectively. (B) *cnot1* knockdown (KD) resulted in low phosphorylation levels and protein amounts of both CLOCK and BMAL1. (C) Densitometric analyses were shown of CNOT1, CLOCK and BMAL1 in control and *cnot1* knockdown U2OS cells from three independent experiments. Data are means (±SD). N=3. A two tailed Student’s *t*-test was performed, and significant differences are represented by black asterisks. (**) *P*<0.01. (D and E) Ser to Ala combined mutations in CLOCK at 440, 441, 478, and 479 sites, or in BMAL1 at 42 and 246 sites resulted in compromised phosphorylation promoted by CNOT1 (D) and increased transactivation (E). (F) PKA-Cα inhibited transcriptional activity of CLOCK-BMAL1.

### Both CLOCK and BMAL1 are PKA targets and CNOT1 promotes association between CLOCK or BMAL1 and endogenous PKA

If CLOCK and BMAL1 are targets of PKA, direct or indirect protein interactions would be expected to occur. To confirm these protein interactions, we coexpressed 3HA-PKA-Cα respectively with 3Flag-tagged CLOCK, BMAL1 or CNOT1-C in HEK293T cells. Immunoprecipitation (IP) results showed that both CNOT1 and BMAL1 were bound to PKA (Fig 3A). In the absence of CNOT1, weak interaction between CLOCK and PKA was found, however, addition of CNOT1 increased CLOCK association with PKA (Fig. 3A). If phosphorylation of CLOCK and BMAL1 promoted by CNOT1 is indeed carried out by PKA, then the phospho-PKA substrate antibody should be able to specifically recognize the consensus (R/K)(R/K)X(S*/T*) sites in CLOCK and BMAL1. Supporting our predictions, coexpression of CNOT1 modestly increased the PKA phosphorylated signal of both CLOCK and BMAL1. Furthermore, coexpression of PKA dramatically elevated phosphorylation levels of these two transcriptional activators. In contrast, the PKA dead kinase (K73E and K169E) (46) failed to promote phosphorylation of both CLOCK and BMAL1 in the same procedure of experiments (Fig. 3B). In addition, overexpression of CLOCK, BMAL1 and CNOT1 could pull down endogenous PKA respectively (Fig. 3C), consistent with the IP results. Coexpressing CNOT1 increased the endogenous PKA activity pulled down by over-expressing CLOCK or BMAL1 (Fig. 3D and E). Collectively, these data suggest that the PKA-promoted phosphorylation of both CLOCK and BMAL1 could be regulated by CNOT1 in mammalian cells.

**FIGURE 3.**
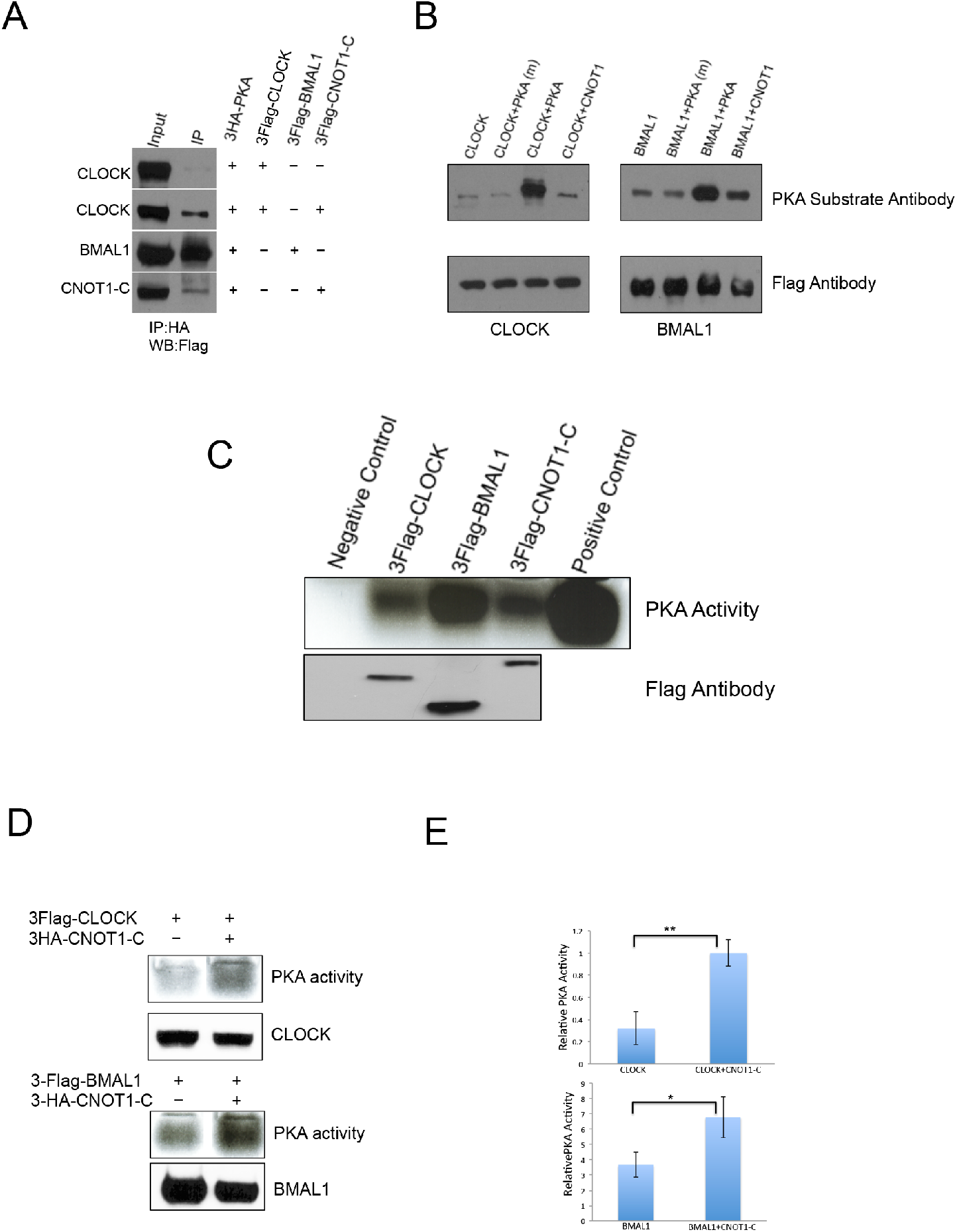
CNOT1 promotes associations between CLOCK or BMAL1 and endogenous PKA. (A) Interactions were shown between PKA and BMAL1, CNOT1, and CNOT1 increased the association between CLOCK and PKA in HEK293T cells in overexpressing conditions. (B) PKA-Cα could, but the PKA dead kinase (K73E and K169E) could not, phosphorylate both CLOCK and BMAL1. (C) Overexpressing Flag-tagged CLOCK, BMAL1 and CNOT1-C could pull down endogenous PKA, respectively. HEK293T cells were transfected by different plasmids and crude protein samples were pull-downed by Anti-Flag Affinity Gel. Negative control is mock transfection. The commercial PKA from the kit acts as a positive control. (D) CNOT1 promoted both CLOCK and BMAL1 binding to endogenous PKA. (E) Densitometric analyses were shown from three independent experiments. Data are means (±SD). N=3. A two tailed Student’s *t*-test was performed, and significant differences are represented by black asterisks. (*) *P*<0.05; (**) *P*<0.01.

### CNOT1 associates with CLOCK in a circadian rhythm and promotes CLOCK phosphorylation in liver

To test whether the same mechanism also works *in vivo*, we used two transgenic mouse lines: one is tetO::*Clock^wt^*-HA (47); another line is called LAP–tTA in which the tTA is expressed under the control of the liver activated protein promoter (48). After breeding, animals expressing CLOCK-HA specifically in liver were generated (49,50). Then HA immuno-precipitation was performed to purify CLOCK in liver (Fig. 4A). The mass spectrometry analysis identified a few endogenous proteins co-purified together with CLOCK-HA, such as BMAL1, DDB1(51), OGT(10) and ZBTB20(52), suggesting constitutively expressed CLOCK-HA play a same role as endogenous CLOCK. Most interestingly, both CNOT1 and CNOT2 associated with CLOCK (Fig. 4B) in the same purification. Both CNOT1 and CLOCK-HA protein levels showed no difference between ZT8 and ZT16 (Fig. 4C and D). The HA antibody could pull down CNOT1 only at ZT8, not at ZT16, indicating there is rhythmic interaction between CLOCK and CNOT1 in liver (Fig. 4C). Consistent with these results, both CLOCK and BMAL1 showed more phosphorylation levels regulated by PKA at ZT8 than those at ZT16 (Fig. 4D), suggesting that PKA can promote clock proteins phosphorylation via CNOT1 in mice.

**FIGURE 4.**
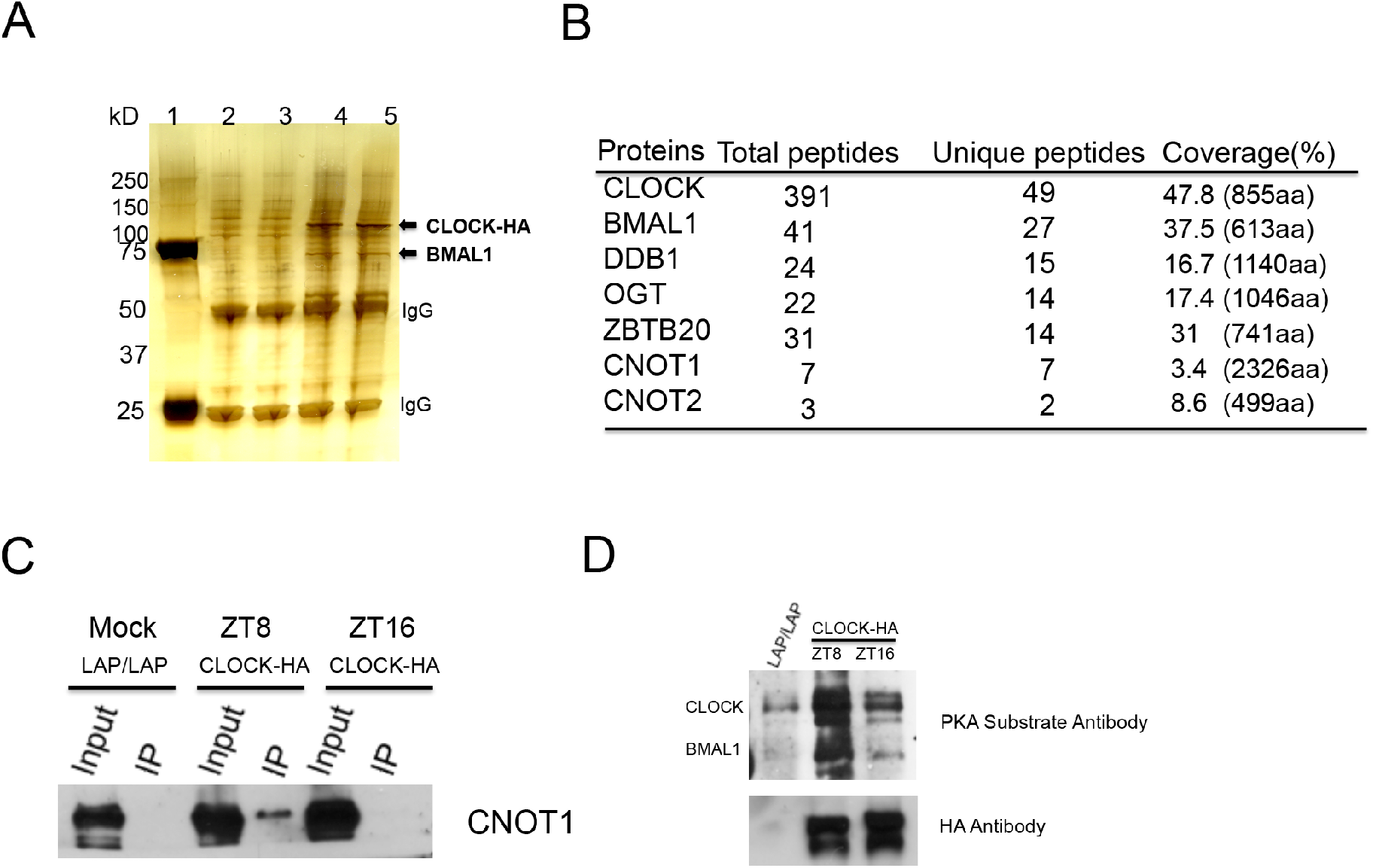
CNOT1 associates with CLOCK BMAL1 and promotes their phosphorylation by PKA in liver. (A) Anti-HA immuno-precipitation samples were shown on a SDS-PAGE gel. Lane1, protein markers; lane 2&3, controls from LAP/LAP mice liver; lane 4&5 from LAP/TETO::CLOCK-HA mice liver. (B) Mass spectrometry results identified a few endogenous proteins co-purified together with CLOCK-HA in liver. (C) Associations between CNOT1 and CLOCK were stronger at ZT8 than ZT16 in liver. (D) Accordingly, PKA phosphorylation levels of both CLOCK and BMAL1 were higher at ZT8 than ZT16 in liver.

### PER2 is also a PKA target

In *Neurospora*, PKA regulates both WC proteins and FRQ phosphorylation (32). We asked whether PKA could function on PER and CRY in mammalian cells. After over-expressing Flag-PER2 and HA-CRY2 separately in HEK293T cells, the PKA substrate antibody only recognized PER2, not CRY2, pulled down by respective IP, suggesting that only PER2 is a PKA target (Fig. 5A). Not surprisingly, both PKA and CNOT1 promoted PER2 phosphorylation (Fig. 5B), and PKA phosphorylation increased stability of the PER2 proteins (Fig. 5C).

**FIGURE 5.**
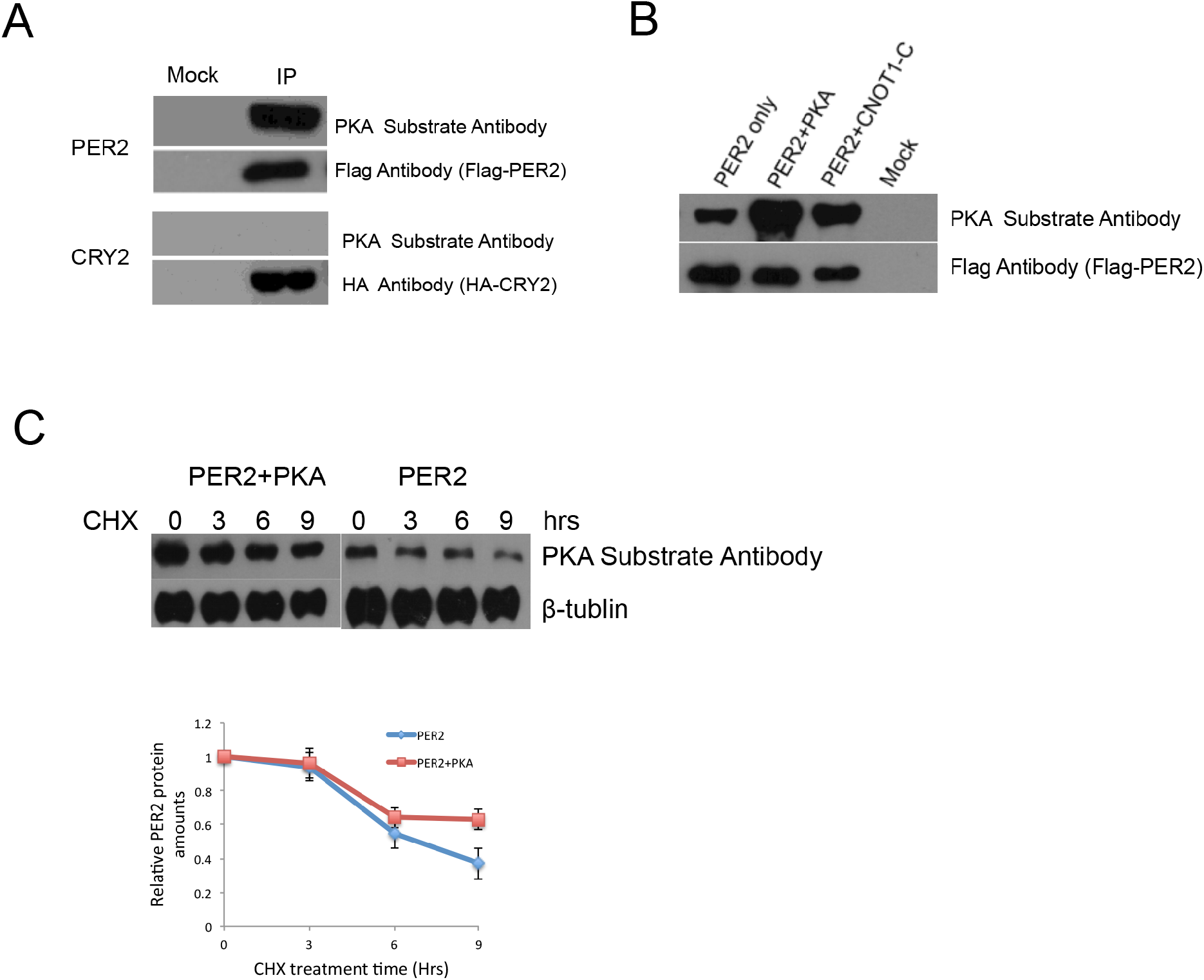
PER2, not CRY2, is also a PKA target, and both PKA and CNOT1 promote PER2 stability. (A) IP results showed that PER2 could be recognized by the Phospho-PKA substrate antibody. (B) Coexpressing transfection results demonstrated that both PKA and CNOT1 increased PER2 protein levels. (C) CHX treatment experiments indicated that PKA phosphorylation increased PER2 protein stability. Densitometric analyses were shown from three independent experiments.

### PKA genetic inhibition changes the period of circadian cycles

Because PKA regulates phosphorylation of CLOCK, BMAL1 and PER2, we predict that changing its activity would affect circadian cycles. In mammals among three PKA catalytic subunits –α, -β and -γ, the *PKA-Cα* gene plays a major role (53). Using U2OS cells bearing the *Bmal1-luc* reporter gene, a 20-nt guide sequence targeting the exon3 of the *PKA-Cα* gene was designed to generate mutants with CRISPR-Cas9 (54) (Fig. 6A). The sequencing data showed that 5-nt base pairs were deleted immediately upstream of the PAM sequence in a mutant developed from a single colony (Fig. 6B). The PKA activity was dramatically lower in this deletion mutant than WT; while the U2OS stabilized cell line over-expressing PKA-Cα showed higher PKA activity than WT (Fig. 6C). Both PKA-Cα over-expressing cells (OX) and deletion mutants (MT) showed dampened rhythmicity after two cycles (Fig. 6D), indicating that an optimal level of PKA activity is important for circadian oscillations and other cellular functions. In addition, the phases between the deletion and over-expressing mutants were modified in an opposite way, the former advanced and the latter delayed for about 2 hours (Fig. 6D). Interestingly, the PKA deletion strain exhibited a longer period of circadian cycles, on the contrary, the PKA over-expressing strain showed a shorter period (Fig. 6E). Taken together, these results indicated that PKA plays an important role in the mammalian circadian feedback loop.

**FIGURE 6.**
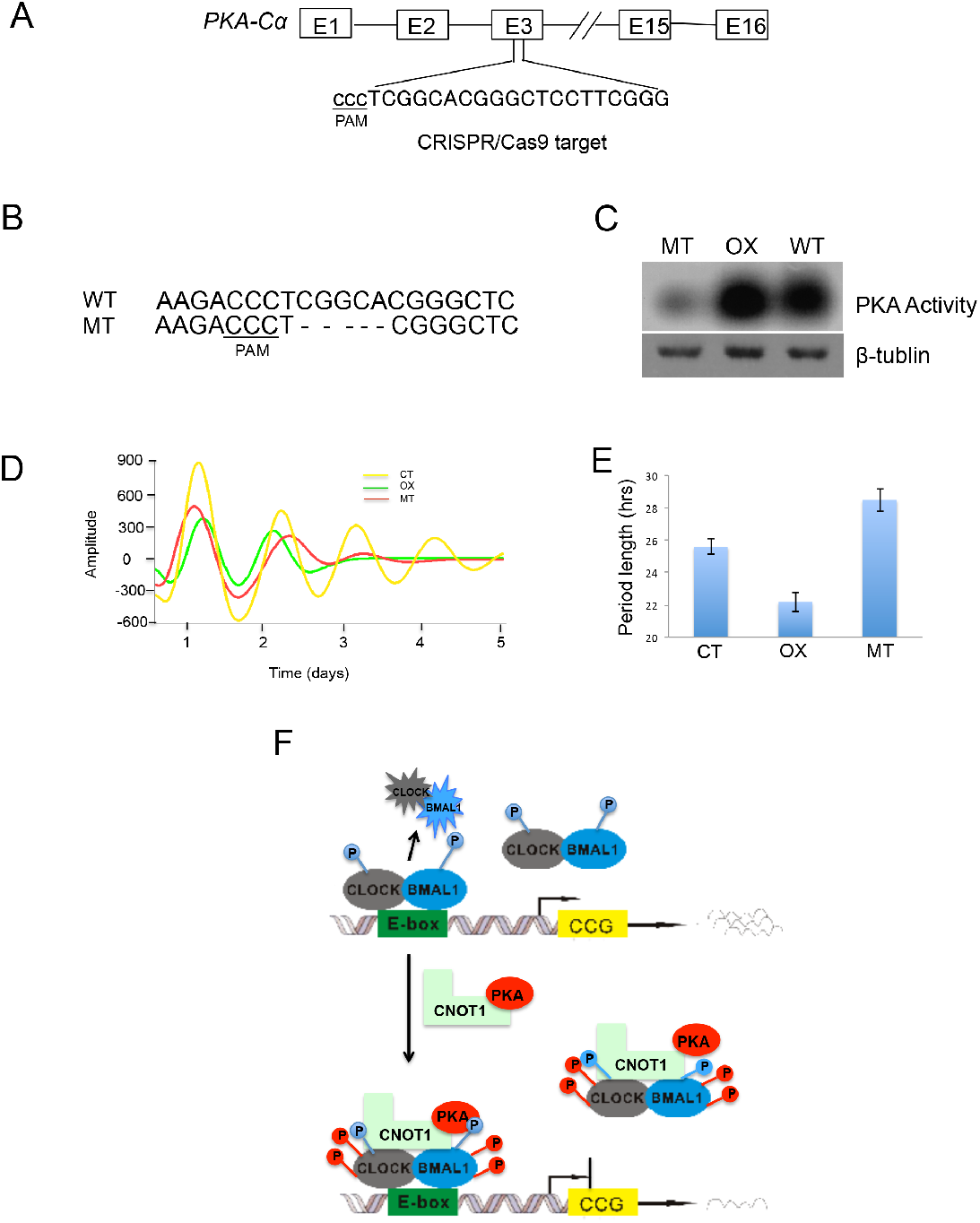
Genetic deletion of *PKA-Cα* affects circadian cycles. (A) The target DNA location by CRISPR-Cas9 was shown in the *PKA-Cα* gene. (B) Sequencing results showed a 5-nt deletion preceding the PAM sequence. (C) The *PKA-Cα* mutant and the PKA overexpressing mutant respectively exhibited lower and higher PKA activity than that of WT. (D) The amplitudes and phases of both PKA overexpression and deletion mutants were altered in the circadian cycles. WT: wild type; OX: PKA overexpression; MT: PKA deletion mutant. (E) Period lengths were calculated based on four repeat experiments. (F) A schematic model of PKA and CNOT1 mediating phosphorylation and transcription of CLOCK-BMAL1 is shown. CLOCK and BMAL1 form a heterodimer, which is phosphorylated and activated by unknown kinases. Binding to the E-box leads to CLOCK-BMAL1 degradation and downstream clock controlled genes (CCG) transcription. CNOT1 bridges PKA and associates with CLOCK and BMAL1, increasing their phosphorylation and protein stability, in turn inhibiting their transactivation.

## Discussion

Both PER and CRY proteins function as essential repressors in the mammalian circadian feedback loop (55). Interestingly, two components unique to the vertebrate genome, CIPC (CLOCK-Interacting Protein, Circadian) (56) and CHRONO/JM129 (57–59), are reported to inhibit CLOCK–BMAL1 independent of CRY. Nevertheless, the polycomb group protein EZH2 is recruited by CLOCK-BMAL1 and represses target genes expression by di- and trimethylation of H3K27 in a CRY-dependent manner (60). Furthermore, another two components of the polycomb repressor complex (PRC2), EED and SUZ12 also interact with CLOCK and BMAL1 to inhibit the *ccl2* gene expression (61). DEC1 and DEC2, two basic helix-loop-helix transcription factors, could inhibit CLOCK-BMAL1 by competing with CLOCK to interact with BMAL1 (5). Here, we show that CNOT1 promotes both CLOCK and BMAl1 phosphorylation through PKA, leading to transcriptional inhibition of this heterodimer, that is different from the action of CRY, PER and other repressors mentioned above.

Among negative regulators of CLOCK and BMAL1, Fukada and colleagues report that CRY2 suppresses and PER2 has no function on the phosphorylation states of both CLOCK and BMAL1 in coexpressed NIH3T3 cells; CIPC promotes phosphorylation and stabilizes both proteins only in the CLOCK-BMAL1 complex (7). Interestingly, coexpression of CNOT1 with CLOCK or BMAL1 alone stimulates their phosphorylation and increases their protein stability, on the contrary, the *cnot1* knockdown decreases both phosphorylation levels and protein amounts in these two proteins (Fig. 2B and 2C), suggesting that CNOT1 regulates the positive elements of the mammalian clock in a unique manner.

Not1 was originally identified as a transcriptional repressor in yeast (37,38), and it inhibits *frq* transcription in *Neurospora* (41). In humans, CNOT1 represses the ligand-dependent transcriptional activation of the oestrogen receptor (62). However, no underlying mechanisms for repression were reported. In this study, we demonstrate that CNOT1 interacts with CLOCK and BMAL1 and inhibits their transcriptional activity via PKA phosphorylation. The S440, S441, S478 and S479 in CLOCK were potential PKA phosphorylation sites, and quadruple Ser-to-Ala mutations resulted in transactivation increase (Fig. 2E). Moreover, there is rhythmic binding between CNOT1 and CLOCK, resulting in more PKA phosphorylation at ZT8 than ZT16 in liver (Fig. F and G). In accordance with these results, the phosphopeptide amount at the CLOCK S440, S441 and S446 oscillates with a peak around ZT8 and a trough around ZT16 in liver (63), indicating that both CLOCK S440 and S441 might be the PKA phosphorylation sites *in vivo*. However, these results are somewhat unexpected because ZT8 is the activation phase for CLOCK-BMAL1 binding to E-box elements of clock genes. One possibility is that more phosphorylated CLOCK and BMAL1 by PKA at ZT8 ensure total protein levels are the same as at ZT16 in the repressive phase. If DNA-binding leads to degradation of transcriptional factors, we would predict that both levels of CLOCK and BMAL1 are lower at ZT8 than ZT16. By contrast, no large differences are shown for these two protein levels during the circadian cycle (7,64,65), although nuclear accumulation of CLOCK and BMAL1 might oscillate (66). This suggests that constant levels of CLOCK and BMAL1 are important to maintain both circadian and non-circadian regulation in mammals. Indeed, both CLOCK and BMAL1 have additional functions in cells. For example, CLOCK up-regulates NF-kB-mediated transcription independent of BMAL1 (67). BMAL1 interacts with the translational machinery in the cytosol and promotes protein synthesis without CLOCK (68).

PKA is reported as a priming kinase for casein kinases to mediate sequential phosphorylation events in the circadian negative feedback loop in *Neurospora* (32). Both WC-1 and WC-2 are regulated by PKA, and PKA phosphorylation promotes protein stability and inhibits transcriptional activity of the WC complex. In line with these results in fungi, we show here that PKA could phosphorylate and, thus, stabilize CLOCK and BMAL1, resulting in inhibition of CLOCK-BMAL1 transactivation. Unexpectedly, our results show that CNOT1 mediates PKA phosphorylation of clock proteins in mammals. With this idea, we could nicely explain why down-regulation of *not1* leads to low levels of both WC-1 and WC-2 and reduced phosphorylation in fungi (41).

Our data indicated that PKA could also target PER2 phosphorylation and enhance its stability, similar to FRQ regulation in fungi (32), PER (29) and TIMELESS accumulation in *Drosophila* (30), suggesting that PKA is a conserved kinase involved in the circadian clock in widely divergent taxa.

Why does PKA phosphorylation increase CLOCK-BMAL1 stability and inhibit its transcriptional activity? One possibility is that hyper-phosphorylated CLOCK/BMAL1 weakens its DNA binding activity; on the contrary, hypo-phosphorylated CLOCK/BMAL1 exhibits higher transcriptional activity as Kamikaze activators (69), which are rapidly degraded once bound to DNA. Supporting this notion, quadruple Ser-to-Ala mutations in CLOCK and double Ser-to-Ala mutations in BMAL1 at S42 and S246 showed increased transactivation activity and decreased protein stability (Fig. 2E). Consistent with our data, with less protein amounts, BMAL1-S42A bound to E-box elements in the promoters of both *Dbp* and *Nr1d1* is substantially higher than WT BMAL1, and mRNA levels of these two genes are enhanced by the BMAL1 mutation in liver (70). Another possibility is that PKA could directly phosphorylate UBE3A at T485 and inhibit UBE3A ubiquitin ligase activity (71). As UBE3A interacts with BMAL1 and regulates its ubiquitination and degradation (14,15), hyper-phosphorylated BMAL1 could not be ubiquitinated by phosphorylated UBE3A, resulting in BMAL1 accumulation together with CLOCK. Third, phosphorylated CLOCK/BMAL1 could be decreased in nuclear subcellular abundance (7). In contrast to the PKA function in this study, the PKC phosphorylation, nevertheless, increases CLOCK-BMAl1 binding to the E-Box of clock genes (27), suggesting that different kinases play distinct roles in the mammalian circadian feedback loop.

In *Drosophila*, increasing PKA activity by inhibition of the PKA regulatory subunit results in arrhythmic circadian locomotor activity (72). As dPER is stabilized by increasing cAMP levels and cAMP-mediated PKA activity, enhanced dPER stability might lengthen circadian periods (29). However, increasing PKA activity caused a shorter period in U2OS cells in this study. As PKA phosphorylation not only stabilized PER2 (Fig. 5C), but also promoted accumulation of both CLOCK and BMAL1, leading to their transcriptional inhibition (Fig. 2E). In addition, it is reported that negative element turnover is uncoupling from circadian period determination (73). It is still not fully understood why up-regulating PKA activity reduces the circadian period. In contrast, genetic inhibition of PKA activity by CRISPR-Cas9 lengthened the period of U2OS cells, highlighting that PKA plays an important role in regulating the circadian period by posttranslational modifications in mammals. In addition, PKA deletion and over-expression resulted in opposite phase changes of circadian cycles, reminiscent of the Ccr4-Not protein complex regulating the phase of the *Neurospora* circadian clock (41).

The Co-IP results showed that there are no significant associations between CLOCK and PKA under the overexpression condition. Addition of expressing CNOT1 obviously promoted their interactions and direct binding between CNOT1 and PKA was detected (Fig. 3A), suggesting CNOT1 might bridge PKA for CLOCK phosphorylation *in vivo*. We also observed interactions between BMAL1 and PKA without heterologous CNOT1 (Fig. 3A). One possibility is that endogenous CNOT1 bridges BMAL1 to PKA. Secondly, it is likely that BMAL1 directly associates with the PKA holoenzyme, which is composed of two regulatory subunits and two catalytic subunits (74); CNOT1 binding to BMAL1 results in the PKA conformation change, which is more accessible for cAMP binding and regulatory subunits releasing. Of note, the cAMP oscillation (75) possibly provides another layer to regulate PKA activity and circadian phosphorylation of clock proteins. Taken together, our data reveal that CNOT1 promotes phosphorylation of mammalian clock proteins via PKA. We propose a possible schematic model to show how PKA and CNOT1 regulate CLOCK-BMAL1 phosphorylation and transcription (Fig. 6F).

What is the nature of *cnot1* transcriptional regulation? The qPCR and RNA-Seq results showed a low amplitude of *cnot1* changes in a circadian cycle in liver (data not shown), consistent with the data shown using Affymetric Mouse Genome arrays (76). Furthermore, the same research reports that there is no obvious oscillation of the *cnot1* gene in human cells (76). In a genome-wide RNAi screen in U2OS cells, *cnot1* knockdown results in a phase delay and low amplitude of clock-controlled luciferase rhythm (77). By performing Mass Spectrometry of samples pulled down together with CLOCK-HA in liver, not surprisingly, we found that a few proteins, such as BMAL1, OGT (O-GlcNAc Transferase) (10,11), ZBTB20 (52), etc., associate with CLOCK and BMAL1; more strikingly, both CNOT1 and CNOT2 are also on the CLOCK-bound list. Future work will focus on other components of the CCR4-Not complex in regulating core clock proteins in mammals.

## Materials and methods

### Cell cultures

HEK293T cells were grown in the Dulbecco’s Modified Eagle Medium (DMEM) supplemented with 10% fetal bovine serum, 100 U ml^−1^ penicillin and 100μg ml^−1^ streptomycin. U2OS cells were grown in the McCoy’s 5A medium with the same supplements as HEK293T cells. Cell cultures were regularly maintained in a 5% CO_2_ incubator. In terms of cells for luminescence recording, they were grown in the DMEM supplemented with 0.035% sodium bicarbonate, 10mM HEPES, 1% fetal bovine serum, 2mM L-glutamine, 1mM sodium pyruvate, 100 U ml^−1^ penicillin, 100μg ml^−1^ streptomycin, 100 μM luciferin and 100nM dexamethasone (Dex).

### Mice

The mice were housed under 12-h light/12-h dark cycle and followed free access to normal chow diet. All animal care and experimental procedures were performed in accordance with University of Texas Southwestern Medical Center IACUC guidelines, approved by the Ethical Review Committee at the University of Southwestern Medical Center and performed under the IACUC-2009-0054 protocol.

### Plasmids

The plasmid carrying the full length of *cnot1*, pT7-EGFP-C1-HsNot1, was a gift from Elisa Izaurralde (Addgene plasmid # 37370) (78). Three fragments of the C-terminal, N-terminal and middle part were cloned base on the pT7-EGFP-C1-HsNot1. In terms of genetic deletion of PKA-Cα, pST1374-NLS-flag-Cas9 (Addgene plasmid # 44758) and pGL3-U6-sgRNA-PKG-puromycin (Addgene plasmid #51133) were gifts from Xingxu Huang (79) (80). *PKA-Cα, mCLOCK, mBMAL1 mPER2* and *mCRY2* were cloned in the 3Flag or 3HA-tagged pcDNA3.1 vector or as otherwise indicated.

### Luciferase assay

HEK293T cells (5 × 10^4^ per well, in 24-well plates) were lysed in 100 μl of lysis buffer after 24 h transfections. Luminescence was measured following the kit protocol from Promega.

#### Real-time luminescence monitoring of circadian oscillations

Luciferase activity was recorded in real time using a 32-channel luminometer (LumiCycle, Actimetrics) that was maintained at 37°C in a walk-in warm room. Stabilized cell lines were synchronized by Dex before luminescence recording.

### PKA assay

PKA assays were performed using the PepTag nonradioactive PKA assay kit (Promega) following the manufacture’s protocol. The reactions were kept for 30 min at 30°C. PKA activity was visualized by agarose gel electrophoresis of a fluorescent PKA substrate. The gels were photographed under UV light.

### Co-immunoprecipitation (IP)

In terms of interactions between CLOCK or BMAL1 and CNOT1, respective 10μg plasmids of 3Flag-CLOCK, 3Flag-BMAL1 and 3HA-CNOT1-C were transfected by the calcium phosphate-based kit (Clontech) in 10cm plates of HEK293T cell cultures. After 24 hours, cells were lysed by the Passive Lysis Buffer (Promega) supplemented with the Protease Inhibitor Cocktail (Sigma). 1ml supernatant (2mg protein) was incubated with 30μl Protein G Sepharose (GE Healthcare) for two hours on a rotator at 4°C. Then, the supernatant was incubated with 30μl anti-FLAG M2 Affinity Gel (Sigma) overnight at the same condition. The Gel was pelleted and washed three times with the same buffer. 30ul loading buffer was added to the Gel and heated for 5 minutes at 100°C. The supernatant was loaded in each protein lane of SDS-PAGE, after electrophoresis, proteins were transferred onto PVDF membrane and Western blot analyses were performed. To analyze the phosphorylation profiles of CLOCK and BMAL1, a ratio of 149:1 acrylamide/bisacrylamide was adopted for SDS-PAGE gels. Otherwise, SDS-PAGE gels with a regular ratio of 37.5:1 acrylamide/bisacrylamide were used.

### Extraction of mouse liver nuclear proteins and immune-precipitation

Liver nuclear extract was performed as described previously (81) (7). 1 ml final supernatant (2mg proteins) was incubated with 30μl Mouse IgG-Agarose for two hours on a rotator at 4°C. Then, the supernatant was incubated with 30μl Monoclonal Anti-HA Agarose (Sigma) overnight at the same condition. After washing three times with PBS, 30ul loading buffer was added to the gel and heated for 5 minutes at 100°C. Following the electrophoresis and membrane transferring, Western blot analyses were performed using different antibodies.

### Mass spectrometry analysis

Protein samples pulled down by HA-Agarose were run 10mm into the top of a SDS/PAGE gel, stained with Coomassie Blue, and excised. Overnight digestion with trypsin was performed after reduction and alkylation with DTT and iodoacetamide. The resulting samples were analyzed by tandem MS using a QExactive mass spectrometer (Thermo Electron) coupled to an Ultimate 3000 RSLC-Nano liquid chromatography system (Dionex). The detailed procedure and raw data analysis are based on the method published before (82).

### Immunoblotting assay

PVDF membranes carrying proteins were incubated in the TBST buffer with 5% nonfat dry milk (BIO-RAD, Blotting-Grade Blocker, catalogue No. 170-6404) overnight with corresponding antibodies at 4°C on a shaker. If second antibodies are used, then one hour incubation is needed at room temperature. The protein bands were developed by the ECL buffer (GE Healthcare, ECL Prime Western Blotting Detection Reagent, RPN2232), and visualized on films. Primary antibodies in this study were listed as followed: FLAG-HRP (Sigma, M2, catalogue No. A8592); HA-Tag (6E2) Mouse mAb (HRP Conjugate) (Cell Signaling, catalogue No. 2999); Phospho-(Ser/Thr) PKA Substrate Antibody (Cell Signaling, catalogue No. 9621); CNOT1 antibody (Sigma, HPA046577); pAb anti-BMAL1 (NOVUS, catalogue No. NB100-2288).

### Generation of the PKA-Cα deletion and overexpressing mutants

The U2OS cell line bearing the *Bmal-luc* reporter gene (83) was used for *PKA-Cα* genetic deletion by CRISPR-Cas9. A 20-nt DNA sequence preceding the PAM sequence (Fig. 5A) was inserted at the BsmBI site in the pGL3-U6-sgRNA-PKG-Purimycin plasmid. Then two plasmids, one bearing 20-nt target DNA and another one, pST1374 bearing the Cas9 gene, were co-transfected into U2OS cells. After 24 hours transfection, 2 μg/ml puromycin was added into medium for selection. Genomic DNA was extracted from mutants developed from single colonies, and PCR products were cloned for sequencing. In terms of the PKA-Cα overexpressing cells, the plasmid pcDNA3.1.PKA-Cα was transfected into the same U2OS cells, which were selected with 200 μg/ml G418.

### Generation of *cnot1* knockdown mutants

The U2OS cell line was used for *cnot1* knockdown. Two plasmids bearing *cnot1* shRNA are pGIPZ ShRNAmir Lentiviral (Cat #: RHS4430-99292147 and RHS4430-99292155). The plasmid of Non-silencing Verified Negative Control (Cat # RHS4346) was used for the technical manual provided by GE Healthcare. negative control. The protocol was based on the technical manual provided by GE Healthcare.

## Acknowledgments

We thank Drs. Yi Liu, Shin Yamazaki, John Humphreys and Radha Akella (University of Texas Southwestern Medical Center at Dallas) for critical comments regarding the manuscript. We thank Lisa Thomas for providing transgenic animals expressing LAP::tTA and tetO::CLOCK-HA. The plasmid bearing the *mPer-1-Luc* reporter gene was a gift from Dr. Sato Honma. The plasmids for *cnot1* knockdown are from Dr. Jerry Shay’s laboratory in University of Texas Southwestern Medical Center. The CLOCK antibody is from Dr. Choogon Lee in Florida State University.

## FOOTNOTES

This work was supported by National Natural Science Foundation of China Grant 31571206 and 31271281 (to G.H.). J. S. T. is an investigator in the Howard Hughes Medical Institute.

